# An endogenous retrovirus insertion disrupting bovine *ALKBH8* causes a failure-to-thrive syndrome with immunodeficiency associated with juvenile mortality in Brown Swiss cattle

**DOI:** 10.64898/2026.07.09.737535

**Authors:** Sidonia Glatthard, Naveen K. Kadri, Franz R. Seefried, Laura R. Voitl, Bettina A. Weber, Hermann Schwarzenbacher, Seraina L. Meister, Corinne Gurtner, John F. O’Grady, Meret Osbahr, Alexander S. Leonard, Mireille Meylan, Hubert Pausch, Cord Drögemüller, Joana Jacinto

**Author notes:** These authors contributed equally (Sidonia Glatthard, Naveen K. Kadri). Corresponding author: Joana Jacinto, Bremgartenstrasse 109a, 3012 Bern, Switzerland.

## Abstract

The Brown Swiss (**BS**) cattle breed is one of the major Swiss dairy breeds. Intensive selection and the widespread use of few elite sires in artificial insemination have increased inbreeding and the occurrence of deleterious recessive alleles in the homozygous state. Analyzing life trajectories in large, genotyped cohorts can identify hidden recessive disorders that are difficult to detect using traditional case-control association testing. Long-read DNA sequencing enables precise detection of causal alleles, including structural variants. This study aimed to (1) identify cryptic recessive loci affecting rearing performance in Swiss BS cattle, (2) evaluate their impact on survival, (3) characterize the associated phenotype, (4) identify the causal variant using long-read whole-genome sequencing, and (5) assess its functional impact. Using Homozygous Haplotype Enrichment/Depletion (**HHED**) mapping, we identified a risk haplotype (**BH39**) on chromosome 15 spanning from 16,276,819 bp to 16,446,984 bp that was associated with increased juvenile mortality within the first 180 days of life when present in the homozygous state. The BH39 occurred at a frequency of approximately 4.5% in Swiss BS cattle and 5.3% in German and Austrian BS cattle, and homozygous carriers exhibited a significantly reduced first-year survival rate. Five females homozygous for BH39 underwent clinical examination. They all showed recurrent respiratory disease, impaired growth, poor body condition, rough hair coat, and brown-discolored teeth. Pathological examination revealed bronchopneumonia and eosinophilic enteritis.

Clinicopathological findings indicated failure to thrive and immunodeficiency. Long-read WGS of two BH39 homozygous calves revealed a private homozygous coding variant that was in high linkage disequilibrium with BH39. The identified structural variant was an insertion of a large transposable element (10.4 kb ERVK[2-1-LTR]) into the third exon of *ALKBH8* (NM_001080341.2 c.267_268indel). Full-length RNA sequencing of cerebellum and liver from a homozygous calf revealed that the endogenous retrovirus (**ERV**) insertion introduces a cryptic transcription termination signal, truncating *ALKBH8* mRNA. This study demonstrates that exploring population-scale genomic data and mining thousands of life-history records, followed by veterinary follow-up evaluations and molecular genetic analyses, provides an effective strategy for identifying cryptic recessive disorders that shorten the lifespan of cattle. The findings provide strong evidence that the ERV insertion into the coding sequence of *ALKBH8* represents a loss-of-function variant that causes a previously undescribed recessive disorder that results in increased rearing loss.

**Interpretive summary:** We identified a recessive disorder in Brown Swiss cattle that causes retarded growth, recurrent infections, immunodeficiency, and increased mortality during the first year of life. Using population-scale genomic data, clinical investigations, and long-read sequencing, we linked the disorder to an exonic transposable element insertion disrupting *ALKBH8*. The identification of the causal variant now enables direct genetic testing and the implementation of genome-based mating strategies to avoid carrier-by-carrier matings and, consequently, prevent the birth of affected homozygous offspring. We demonstrate the utility of integrating large-scale breeding records, veterinary phenotyping, and advanced genomics to identify hidden defects affecting livestock health and productivity.

## INTRODUCTION

Brown Swiss (**BS**) cattle represent one of the most important dairy breeds in Switzerland, appreciated for their high milk quality, high feed efficiency, and adaptability to alpine farming conditions (De Marchi et al., 2007; Bosch et al., 2025; Braunvieh Schweiz, 2026). Intense selection and the widespread use of a small number of elite breeding sires have increased inbreeding levels, resulting in the phenotypic manifestation of deleterious recessive alleles (Hiltpold et al., 2020; Häfliger et al., 2021b; He et al., 2026). Recessive conditions associated with embryonic losses, unspecific phenotypes, or disease onset outside the perinatal period are difficult to detect with commonly applied monitoring programs (Besnard et al., 2024).

Recent studies have demonstrated the value of integrating routine longitudinal phenotype data with extensive genomic information to reveal hidden deleterious alleles (Häfliger et al., 2021a, 2022; Reynolds et al., 2021). Besnard et al. (2024) introduced a novel genome-wide Homozygous Haplotype Enrichment/Depletion (**HHED**) mapping approach, scanning haplotypes for segments that occur disproportionately often in the homozygous state in animals that die young while being largely absent in healthy adult cows. Such a pattern is consistent with recessive inheritance of life-shortening haplotypes that can escape detection through traditional case/control association analyses. A notable example of the application of this approach was the discovery of Bovine Lymphocyte Intestinal Retention Defect (**BLIRD**), a recessive monogenic disorder in Holstein cattle caused by a missense variant in the *ITGB7* (Besnard et al., 2024). Homozygous animals with BLIRD exhibit impaired localization of CD4 T lymphocytes in the intestinal lamina propria, resulting in retarded growth, altered hematological parameters, and an increased risk of juvenile mortality (Dutheil et al., 2025). Due to variable expressivity, incomplete penetrance, and the absence of a pathognomonic clinical phenotype, BLIRD remained undetected in the population for an extended period, illustrating how HHED mapping enables the identification of cryptic, life-shortening Mendelian disorders (Besnard et al., 2024).

Once such candidate regions have been identified at the population level, detailed molecular genetic characterization of the underlying causal variants is indicated. Advances in DNA sequencing technology have transformed the ability to characterize the molecular basis of such defects (Moresco et al., 2013; van Dijk et al., 2018; Espinosa et al., 2024). Third generation long-read sequencing platforms, including Pacific Biosciences (**PacBio**) and Oxford Nanopore Technologies (**ONT**), now enable highly contiguous genome assemblies and precise detection of small and structural variants (Moresco et al., 2013; van Dijk et al., 2018; Espinosa et al., 2024).

To date, no HHED mapping approach has been applied to systematically identify cryptic recessive alleles associated with rearing losses in BS cattle. Therefore, the aims of this study were to (1) apply a HHED mapping approach to identify cryptic recessive loci affecting rearing performance in Swiss BS cattle (≤180 days) by leveraging large-scale life history and pedigree records, (2) evaluate risk haplotype occurrence and effect on animal survival, (3) comprehensively characterize the associated clinical and pathological phenotypes in homozygous animals, (4) identify potential causal variants using long-read whole genome sequencing (**WGS**), and (5) assess the functional impact of the identified candidate variant.

## MATERIALS AND METHODS

### Datasets

Microarray-derived genotypes of 74,225 animals from the BS and Original Braunvieh (**OB**) populations were considered. Animals were genotyped using ten different single nucleotide variant (**SNV**) arrays comprising between 20,000 and 777,000 markers. The positions of the SNV were determined based on the ARS-UCD1.2 bovine reference genome (BosTau9) (Rosen et al., 2020a; NCBI Bos Taurus Annotation Release 106, 2025). Quality control of the array data was performed using PLINK v1.9 (Purcell et al., 2007) separately for each breed-array combination; samples and autosomal SNVs with >10% missing genotypes were excluded. The genotypes were subsequently imputed to the WGS-level in a two-step procedure using Beagle v5.4 (Browning et al., 2018). First, medium-density array genotypes were imputed to high-density (**HD**) level (Illumina BovineHD BeadChip) using a reference panel consisting of 1,183 BS and OB animals.

Second, genotypes for 26,265,006 sequence variants were imputed into the partially imputed HD genotypes using a reference panel of 607 in-house sequenced animals as described previously (Lloret-Villas et al., 2021; Watson et al., 2026). A total of 58,871 animals with a pedigree-derived proportion of BS ancestry of at least 85% were retained for this study. These animals, representing purebred BS animals, had genotypes for 24,068,946 SNVs. Functional consequences of variants were predicted with the Variant Effect Predictor (**VEP**) software (McLaren et al., 2016a) using cache files obtained from the Ensembl (version 104) annotation of the bovine reference genome.

### Genome-wide HHED mapping

Life-history records were available for a subset (N = 56,766) of the partially imputed BS cohort. This dataset included 1,522 animals (cases) that died on farm during the rearing period (between birth and 360 days of age). It also included 17,305 animals (controls) that survived the rearing period and occurred as parents of at least one offspring.

Cases and controls were retained only if they had two genotyped ancestors (either both parents or one parent and a grandparent from the ungenotyped parent’s lineage). This filtering step reduced the number of cases and controls to 1,231 and 12,952, respectively. Cases were assigned to 12 age-at-death categories (before 30 days, before 60 days, before 90 days, and so forth in cumulative 30-day increments up to 360 days). These categories contained between 237 (age-at-death <30 days) and 1,231 (age-at-death <360 days) animals (Table 1).

**Table 1.** Number of Brown Swiss animals included in different cumulative 30-day age-at-death categories considered as case group in homozygous haplotype enrichment depletion mapping.

| **Age categories (in days)** | **Number of animals in case group** |
|---|---|
| 30 | 237 |
| 60 | 473 |
| 90 | 640 |
| 120 | 743 |
| 150 | 824 |
| 180 | 891 |
| 210 | 947 |
| 240 | 1004 |
| 270 | 1067 |
| 300 | 1125 |
| 330 | 1171 |
| 360 | 1231 |

HHED mapping was performed as previously described (Besnard et al., 2024) to identify haplotypes that were enriched in the homozygous state in cases and concurrently depleted in the homozygous state in controls. This analysis was carried out separately in the 12 age-at-death categories including between 237 and 1,231 cases and 12,952 controls.

Haplotypes were constructed from high-density imputed SNV genotypes using sliding windows of 30 consecutive SNVs with 10-SNV overlaps. For each haplotype, we counted the observed number of homozygotes (N_obs_) separately in cases and controls and estimated the expected number (of homozygotes) (N_exp_) based on within-family transmission probabilities from the two genotyped ancestors (i.e., using their haplotype genotypes).

For haplotypes with N_obs_ > 10 in cases, we computed the enrichment/depletion score, En = (N_obs_ – N_exp_) / N_exp_. Following Besnard et al (2024), haplotypes exhibiting a strong enrichment in cases (En ≥ 0.25) and a strong depletion in controls (En ≤ –0.25) were considered associated with juvenile mortality. The haplotype with the strongest enrichment score in cases was considered as the core haplotype in windows where multiple haplotypes exceeded our enrichment/depletion threshold.

### Survival analysis

The identified risk haplotype (**BH39**) was monitored in 58,833 BS animals for which array-derived genotypes were included in the routine genotype preparation and phasing workflows applied by Qualitas AG to prepare data for genomic prediction. The analysis was conducted in R version 3.6.2 using the packages dplyr (Hadley Wickham; Romain François;

Lionel Henry; Kirill Müller; Davis Vaughan, 2026), data.table (Tyson Barrett; Matt Dowle; Arun Srinivasan; Jan Gorecki; Michael Chirico; Toby Hocking; Benjamin Schwendinger; Ivan Krylov, 2026), ggplot2 (Hadley Wickham, 2016), and survival (Terry M. Therneau, 2026). Survival analysis was performed using the Kaplan–Meier estimator (survfit from the survival package), modeling survival time as a function of haplotype genotype classes.

Animals without a recorded death date were treated as right-censored observations. Survival probabilities and confidence intervals were extracted over a defined time range and visualized, showing survival curves per haplotype class. The survival rate of homozygous and heterozygous BH39 carriers was compared with the survival rate of non-carriers separately for males and females. The same analysis was conducted in a dataset consisting of 127,055 BS animals from the Austrian and German populations. Austrian and German population share a joint routine genetic evaluation system.

### Animals and clinical examination

Five animals identified as homozygous for the associated haplotype of the chromosome (chr) 15 region were subjected to clinical examination (cases 1-5). Lung ultrasonography was performed in three animals (cases 1-3) using a 7.5Mz linear probe. Pneumonia severity was scored using a standardized ultrasonographic scoring system (score 1–5), based on the presence of comet-tail artifacts, lobular consolidation, and the number of lung lobes affected by lobar consolidation (Ollivett and Buczinski, 2016). One animal (case 5) underwent a gynecological examination and rectal gynecological ultrasonography.

The animals examined were selected based on being alive and their owners’ consent to the inclusion in the study and clinical examination. The cohort comprised four female calves (cases 1-4) aged with 108 ± 42 days and one 621-day old heifer (case 5).

All animals originated from bovine viral diarrhea virus free herds.

### Blood analyses and coprology

Blood samples were collected from cases 1 – 5. Complete blood counts (**CBC**) were performed for cases 1–4, while serum biochemical analyses were carried out for all five cases. Fecal samples were collected from cases 1–4 and examined using standard flotation techniques to assess the gastrointestinal parasite burden.

### Pathological Examination

Post-mortem examinations were performed on cases 1 and 2 that were euthanized due to poor general condition and failure to thrive. Necropsy was carried out according to standard procedures for bovine pathology, with systematic macroscopic evaluation of all major organ systems. Tissue samples from multiple organs, including lung, liver, gastrointestinal tract, spleen, kidney, heart, and lymph nodes, were collected and fixed in 4% buffered formalin. Formalin-fixed tissues were trimmed, embedded in paraffin, and hematoxylin and eosin (**HE**) stained histological tissue sections were prepared. Slides were then assessed under the light microscope.

### Pedigree data collection and analysis

Pedigree analyses were conducted on the five clinically examined homozygous (cases 1-5) and 38 heterozygous animals carrying the identified haplotype on chr15. Pedigree data were obtained from the BrunaNet database (Braunvieh Schweiz). Common ancestors among all animals were manually identified. An initial overview pedigree for the five homozygous animals was generated using Pedixplorer (Le Nézet et al., 2025). This pedigree was then manually simplified and subsequently expanded to include confirmed heterozygous carrier animals of the haplotype on chr15.

### Long-read WGS and variant calling

High molecular weight DNA was isolated from EDTA blood samples of two homozygous risk haplotype carriers using the HMW Nanobind kit (Pacific Biosciences, Menlo Park, CA, USA). Two HIFI SMRTbell libraries were constructed, equimolarly pooled and run on two SMRT cells 25 M on a PacBio Revio instrument at the Next Generation Sequencing Platform at the University of Bern. A total of 121.4 (case 1) and 115.4 gigabases (case 2) were generated from 6.6 (case 1) and 7.0 (case 2) million HiFi reads respectively.

The reads had a mean length of 18413.4 (case 1) and 16566.7 (case 2) bases and a median length of 17459 (case 1) and 15770 (case 2) bases, respectively with an N50 of 18897 (case 1) and 17156 (case 2) bases, respectively.

The HiFi reads were aligned to the ARS-UCD1.2 reference genome (Rosen et al., 2020b) available on NCBI using pbmm2 (PacificBiosciences/pbmm2, 2025), with the parameters –preset HIFI. Structural variants (**SVs**) and SNVs were called from the alignments using Sniffles2 2.6.2 (Smolka et al., 2024) and DeepVariant 1.6.1 (Poplin et al., 2018) respectively. The Sniffles2 workflow included individual variant calling per sample, followed by a built-in joint variant calling step. Finally, the SVs were annotated using VEP 113.0 (McLaren et al., 2016b).

SNV calling was first carried out on a per-sample basis using DeepVariant 1.6.1, with model type set to PACBIO. GLnexus 1.4.1 (Lin et al., 2018) was then used to merge the called variants, with the config set to DeepVariantWGS (Yun et al., 2020). Annotation of the SNV vcf file was done using snpEff 5.0c (Cingolani et al., 2012). Our control cohort included 84 animals of predominant BS ancestry that had previously been sequenced with PacBio Hi-Fi reads (Mapel et al., 2026) as part of the ongoing sequencing projects (European Nucleotide Archive Project accession number PRJEB46995).

### Targeted genotyping of the 10.4 kb insertion

The 10.4 kb insertion was genotyped using the following three primers: two flanking primers, forward (5′-TCATAGGCTGTCTTGGATTCTTC-3′) and reverse (5′-CAATGGTGGTTTGGGTAATCAAG-3′), and a third primer (5′-GAGATGGTGGTAGGGGACAG-3′) located in the inserted sequence. This allowed for the simultaneous PCR amplification of the wild-type (reference) allele (150 bp) and the mutant (alternate) allele (233 bp) from genomic DNA using AmpliTaq Gold 360 Master Mix (Thermo Fisher Scientific, Waltham, MA, USA), followed by automated capillary electrophoresis using a Fragment Analyzer (Advanced Analytical Technologies, Orangeburg, NY, USA).

### RNA extraction and sequencing

Cerebellum and liver tissues were collected at necropsy from case 2, immediately stored in RNAlater, incubated at +4°C and then stored at –80°C. Total RNA was extracted with RNeasy Mini kit (Qiagen, Steinhausen, Switzerland), following the manufacturer’s protocol. In brief, frozen tissues were homogenized with a MagNA Lyser (Roche) at 6000^rpm for 50s (x2) in RTL Plus Buffer (Qiagen, Steinhausen, Switzerland) with Dithiothreitol. Total RNA purity was assessed with a NanoDrop One (Thermo Fisher Scientific, Reinach, Switzerland), and RNA integrity number (**RIN**) inferred from a Bioanalyzer RNA 6000 Nano assay (Agilent Technologies, Basel, Switzerland). Concentration was estimated with a Qubit 4.0 fluorometer (Thermo Fisher Scientific, Reinach, Switzerland), prior to submission for sequencing.

For each sample, approximately 400ng of total RNA (RIN >7.0) was used to generate barcoded libraries with the Kinnex full-length RNA kit (Pacific Biosciences, Menlo Park, CA, USA), following the manufacturer’s protocol. Libraries were then pooled and sequenced on a 24h movie, 25M SMRT Cell on a PacBio Revio instrument (Pacific Biosciences, Menlo Park, CA, USA).

Sequenced reads were processed using the PacBio IsoSeq workflow (v. 4.3.0; https://isoseq.how). Briefly, raw Kinnex Hi-Fi reads for multiplexed samples on each SMRT Cell were first segmented using skera (v. 1.4.0) by specifying default Kinnex full-length RNA adapters (https://downloads.pacbcloud.com/public/dataset/Kinnex-full-length-RNA/REF-MAS_adapters/MAS-Seq_Adapter_v3/mas8_primers.fasta). Segmented reads from each SMRT Cell were then demultiplexed with lima (v. 2.13.0) using the --isoseq flag and default Kinnex full-length RNA specific cDNA barcode primers (https://downloads.pacbcloud.com/public/dataset/Kinnex-full-length-RNA/REF-primers/IsoSeq_v2_primers_12.fasta). Demultiplexed reads for each sample were then reoriented in the 5’ – 3’ direction and filtered to retain only full-length non-concatemer (**FLNC**) reads with poly-A tails using isoseq refine (v. 4.3.0). These filtered FLNC reads were then aligned to ARS-UCD 2.0 with pbmm2 (v. 1.17.0) using the --preset ISOSEQ parameter. Aligned reads were then visualized using the Integrated Genomics Viewer (**IGV**; v. 2.19.7)(Robinson et al., 2011).

## RESULTS

### A haplotype on chromosome 15 increases juvenile mortality in Brown Swiss cattle

Genome-wide HHED mapping identified a genomic region on chr 15 associated with rearing losses in the first year of life. This region was identified for the <180 days age-at-death category indicating that homozygosity for the associated haplotype increases juvenile mortality. Associated haplotypes were identified in the genomic region from 16,276,819 to 16,818,987 bp, exhibiting enrichment and depletion scores of > 3.7 in cases and < –0.3 in controls, respectively, across the later age-at-death categories (Figure 1a). The strongest signal was observed for the category including calves dying before 180 days of age (En = 4.33 in cases; En = –0.82 in controls). The core haplotype showing the highest enrichment in the homozygous state in cases spanned a genomic region from 16,276,819 to 16,446,984 bp. This haplotype segregated at a frequency of 3.71% in the BS cohort used for the HHED mapping.

**Figure 1.**
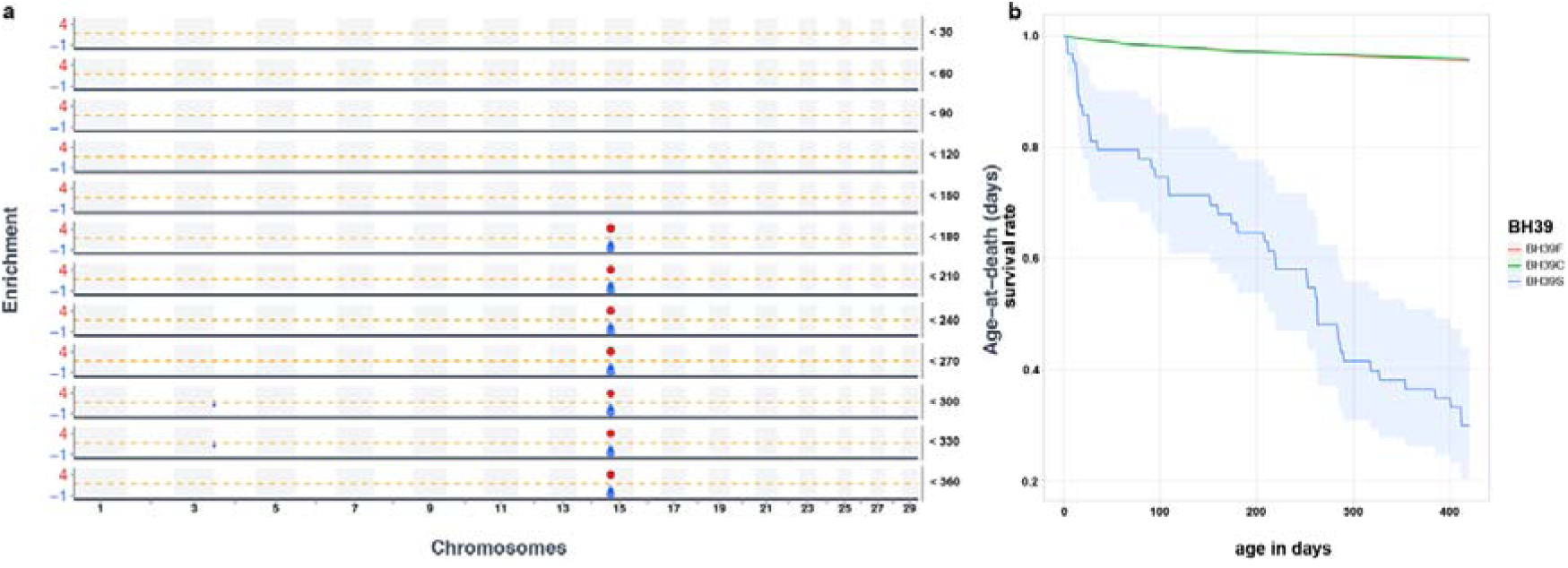
Genome-wide homozygous haplotype enrichment/depletion mapping and juvenile mortality survival analysis in Brown Swiss cattle. a) Genome-wide homozygous haplotype enrichment/depletion mapping and juvenile mortality in Brown Swiss cattle. Enrichment of haplotype homozygosity in cases and controls is shown across age-at-death categories. The results are plotted only for haplotypes exhibiting En >0.25 in cases (red) and concurrent depletion (En < −0.25) in controls (blue). Note the significant risk haplotype designated as BH39 on chromosome 15. b) Female survival rate estimated in 58,833 Swiss BS animals. The blue line represents the survival rate of homozygous BH39 carriers. Blue shaded area is the confidence interval. Survival rate of heterozygous BH39 carriers (green) is indistinguishable from the survival rate of non-carriers (red).

The identified risk haplotype designated as BH39 was integrated into the Qualitas AG routine genotype preparation and phasing workflows applied to prepare data for genomic prediction (Supporting file S1). This allowed the identification of heterozygous and homozygous haplotype carriers among 58,833 routinely genotyped Swiss BS animals. In this cohort, the haplotype segregated at a frequency of 4.54%, with 5,240 carriers and 52 homozygotes (Table 2). Survival rate was analyzed for all three diplotypes of BH39. The survival rate of heterozygous haplotype carriers did not differ from the survival rate of non-carriers. However, the 1-year survival rate was substantially reduced for calves carrying two copies of BH39 (Figure 1b). The observed number of BH39 homozygotes was significantly lower than expected. Usually, Swiss BS calves are genotyped very early in life, before the age of 70 days. Hence, the significant deviation from Hardy-Weinberg proportion (p = 4.25 **×** 10^-11^) might result from a farmer-related exclusion for genotyping at very young age due to reduced health performance or indicate an increased perinatal mortality.

**Table 2.** Occurrence of the BH39 haplotype in Brown Swiss cattle.

| **Cohort** | **ref/ref** | **ref/var** | **var/var** | **Haplotype frequency** |
|---|---|---|---|---|
| Switzerland | 68764 | 5409 | 52 | 3.71% |
| Germany/Austria | 113810 | 13083 | 162 | 5.28% |

The BH39 haplotype segregated at a slightly higher frequency of 5.28% in 127,055 BS animals that were genotyped in the Austrian and German BS population (Table 2). Assuming Hardy-Weinberg proportions, 354 homozygous BH39 carriers would be expected but we observed only 162 (p = 5.32 **×** 10^-27^) in this population. Consistent with observations in the Swiss population, the first-year survival rate of homozygous BH39 carriers was substantially reduced in this cohort (Supporting File S2).

### Failure to thrive and indication of immunodeficiency in BH39 homozygous BS cattle

Amongst the routinely genotyped animals, we identified five living homozygous BH39 females. According to the animal owners, two animals (cases 1 and 3) had a history of recurrent health problems, including repeated episodes of pneumonia, omphalitis, and impaired growth. Overdue birth was reported for four of the five affected animals (Table 3). Case 4 had undergone a laparotomy with cecotomy three weeks prior to the examination. In contrast, the heifer (case 5) had reportedly developed normally; however, fertility problems were reported.

**Table 3.** History and main clinical findings in five BH39-homozygous Brown Swiss females.

| **Case ID** | **Age (in days)** | **History\*** | **General Appearance and Clinical Parameters\*\*** | **Thoracic ultrasonography findings and score\*\*** | **Age at death (in days)** | **Cause of death** |
|---|---|---|---|---|---|---|
| 1 | 99 | Overdue birth. Omphalitis. Episodes of fever, coughing, apathy, reduced appetite, and diarrhea. Slow recovery. Failure to thrive. | Brown discoloration of the teeth. Retarded growth. Cachectic. Thin, dull, and rough hair coat. HR 132/min; RR 44/min; T 39.2 °C. Increased bronchial breath sounds in lungs. | Score 5/5; chronic lobar pneumonia. | 120 | Euthanasia |
| 2 | 92 | Overdue birth. Omphalitis. Failure to thrive. | Brown discoloration of the teeth. Retarded growth. Reduced nutritional status. Thin, dull, and rough hair coat. Omphalitis. Dry cough. HR 80/min; RR 32/min; T 38.4 °C. Increased inspiratory and expiratory sounds in medial lung lobes. | Score 2/5; patchy pneumonia. | 220 | Euthanasia |
| 3 | 62 | Overdue birth. Omphalitis. Episodes of fever, pneumonia, and reduced appetite. Failure to thrive. | Brown discoloration of the teeth. Retarded growth. Reduced nutritional status. Thin, dull, and rough hair coat. Omphalitis. Bilateral purulent nasal discharge. HR 132/min; RR 36/min; T 38.5 °C. Increased inspiratory and expiratory sounds in the left mediodorsal lung lobe. | Score 1/5; pleural irritation and mild interstitial changes. | 109 | Natural death |
| 4 | 177 | Overdue birth. Cecal surgery at 170 days of age due to cecal disorder. | Brown discoloration of the teeth. Retarded growth. Reduced nutritional status. Thin, dull, and rough hair coat. Mild bilateral serous nasal discharge. HR 80/min; RR 44/min; T 39.8 °C. Increased inspiratory and expiratory sounds. | Not performed. | 220 | Natural death |
| 5 | 621 | No report of previous illnesses. Fertility problems. | Brown discoloration of the teeth. Retarded growth. HR 88/min; RR 28/min. | Not performed. | 830 | Slaughter |

\* All information regarding history were based on reports from the respective animal owners.
\*\* On-Farm Use of Ultrasonography for Bovine Respiratory Disease (Ollivett and Buczinski, 2016)
HR = heart rate (Reference Range 70-90 bpm), RR = respiratory rate (Reference Range 15-40 rpm), T = rectal body temperature (Reference Range 38.5-39.5 °C), bpm = heart beats/minute, rpm = respiration/minute

At clinical examination, retarded growth was present in all animals (cases 1-5) (Figure 2a-d). A notable finding was a marked brown discoloration of the teeth (Figure 2e) in all cases. The nutritional status was reduced in all calves (cases 1-4). The four calves (cases 1–4) had a thin, dull, and rough hair coat (Figure 2c). Omphalitis (Figure 2f) was observed in three animals (cases 1-3).

**Figure 2.**
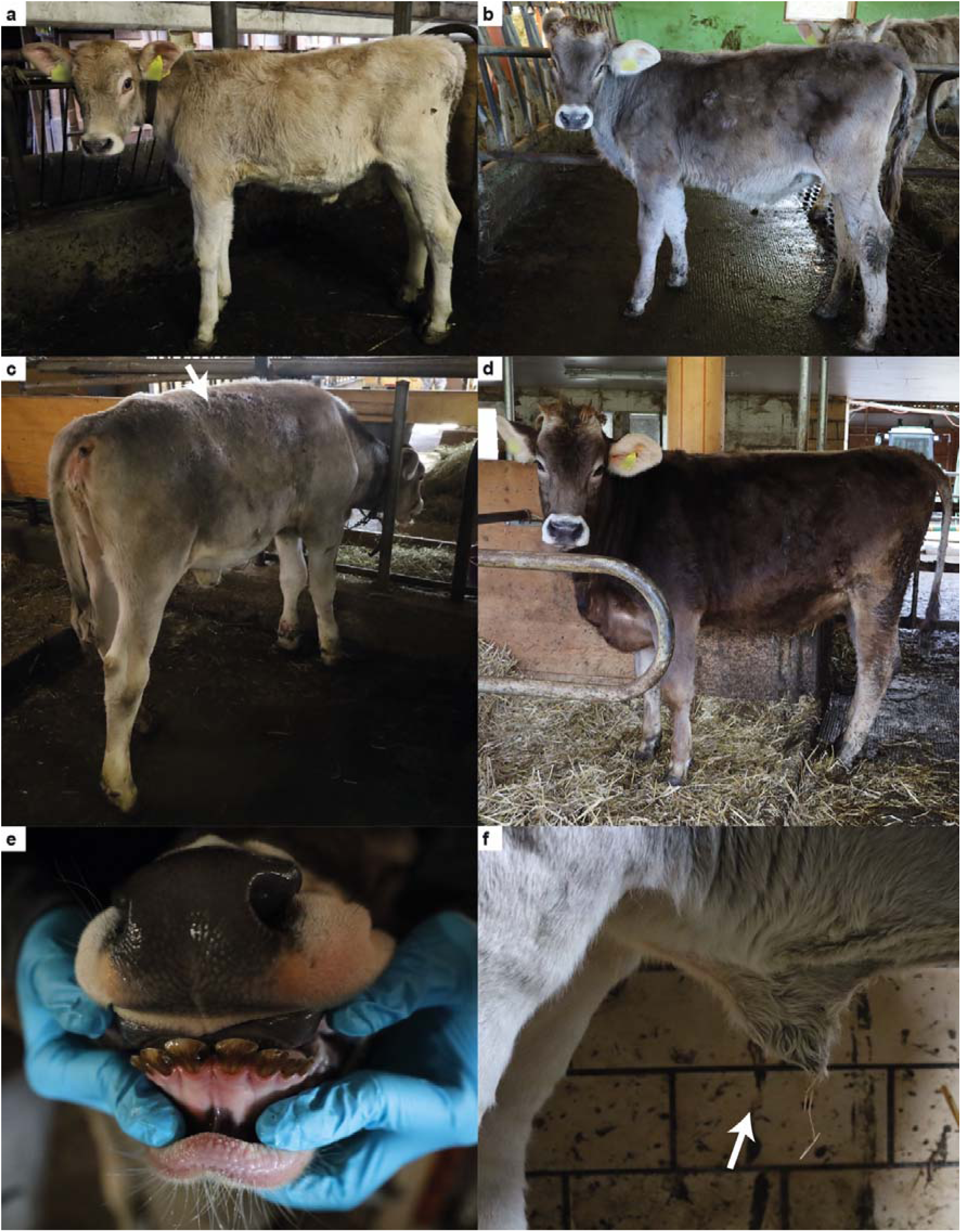
Clinical phenotype in BH39 homozygous Brown Swiss cattle. (a) 120-days-old calf exhibiting emaciation, failure to thrive, dull hair coat, and hypotrichosis (case 1). (b) 219-days-old calf with failure to thrive and reduced nutritional status (case 4). (c) 220-days-old calf showing reduced nutritional status and a dull, woolly, thin hair coat (white arrow) (case 2). (d) 621-days-old heifer showing reduced growth (case 5). (e) Marked brown discoloration of the incisors in a 220-days-old calf (case 2). (f) 109-days-old calf with an enlarged umbilicus (approximately two fingers wide; white arrow) following omphalitis; no increased warmth, pain, or signs of active inflammation were observed at the time of examination (case 3).

Thoracic auscultation revealed abnormal pulmonary sounds in cases 2–4. In three animals (cases 2–4), the cranioventral lung lobes were primarily affected, whereas in case 1 the findings were disseminated throughout the lungs bilaterally.

Thoracic ultrasonography performed in three cases revealed lobar pneumonia in case 1, patchy pneumonia in case 2, and pleural irritation with mild interstitial changes in case 3. All details on clinical examination are summarized in Table 3 and Supporting File S3.

In addition rectal gynecological ultrasonography in case 5 revealed no functional ovarian structures, and the animal was therefore classified as being in anestrus.

Based on the observed clinical findings, the examined animals were suspected to be affected by a multisystemic disorder characterized by recurrent infections early in life, predominantly pneumonia and omphalitis, accompanied by failure to thrive. These repeated disease episodes were consistent with reduced immune competence.

In CBC case 1 had microcytic erythrocytes (31 ± 3 fL, RR 38-54 fL). Case 3 had moderate monocytosis (2.59 x10^9^/L, RR 0.92-1.26 x10^9^/L) and cases 1, 2 and 4 showed moderate monocytopenia (0.46 ± 0.16 x10^9^/L).

In the blood biochemistry, all cases showed elevated asparate aminotransferase (**ASAT**, cases 1-4 83.0 ± 23.2 IU, RR 20.6-48.1 IU; case 5 352 IU, RR 49-151 IU). Cases 1, 2, and 4 showed hypocalcemia (2.22 ± 0.17 mmol/L, RR 2.46-3.66 mmol/L) and increased glutamate dehydrogenase (**GLDH**, 78.0 ± 70.0 IU, RR 3.9-24 IU). Three animals (cases 2, 4, and 5) showed hypoproteinemia (cases 2 and 4 51.0 ± 0.6 g/L, RR 57.0-69.5 g/L; case 5 54.3, RR 65.4-87.3 g/L). Cases 1 and 4 additionally exhibited mild hypoalbuminemia (27.2 ± 2.1 g/L, RR 29.5-37.5 g/L). Case 5 demonstrated marked increased creatine kinase (**CK**, 23276 U/L; RR 71–266 U/L).

The complete blood analysis results, including individual values, units, and reference intervals, are provided in Supporting file S4.

Oocysts of one or more *Eimeria* spp. were detected in all four animals examined (cases 1–4). The oocyst burden was assessed semi-quantitatively using a scoring system ranging from + to +++.

### BH39 homozygous BS cattle have unspecific lung and intestine lesions

The two cases subjected to necropsy (cases 1 and 2) had bronchopneumonia predominantly affecting the cranial lung lobes. In case 1, bronchopneumonia extended to the middle lung lobes (Figure 3a-b). The affected lung areas were firm on palpation and clearly demarcated from the surrounding parenchyma. In case 2, the pulmonary lesions were less extensive and involved approximately 15% of the total lung parenchyma (Figure 3c). Furthermore, both animals showed multifocal dark brown to black discoloration of the teeth (Figure 3d-f). Lymphadenomegaly was observed, affecting the tracheobronchial lymph nodes in case 1 and the mesenteric lymph nodes in case 2.

**Figure 3.**
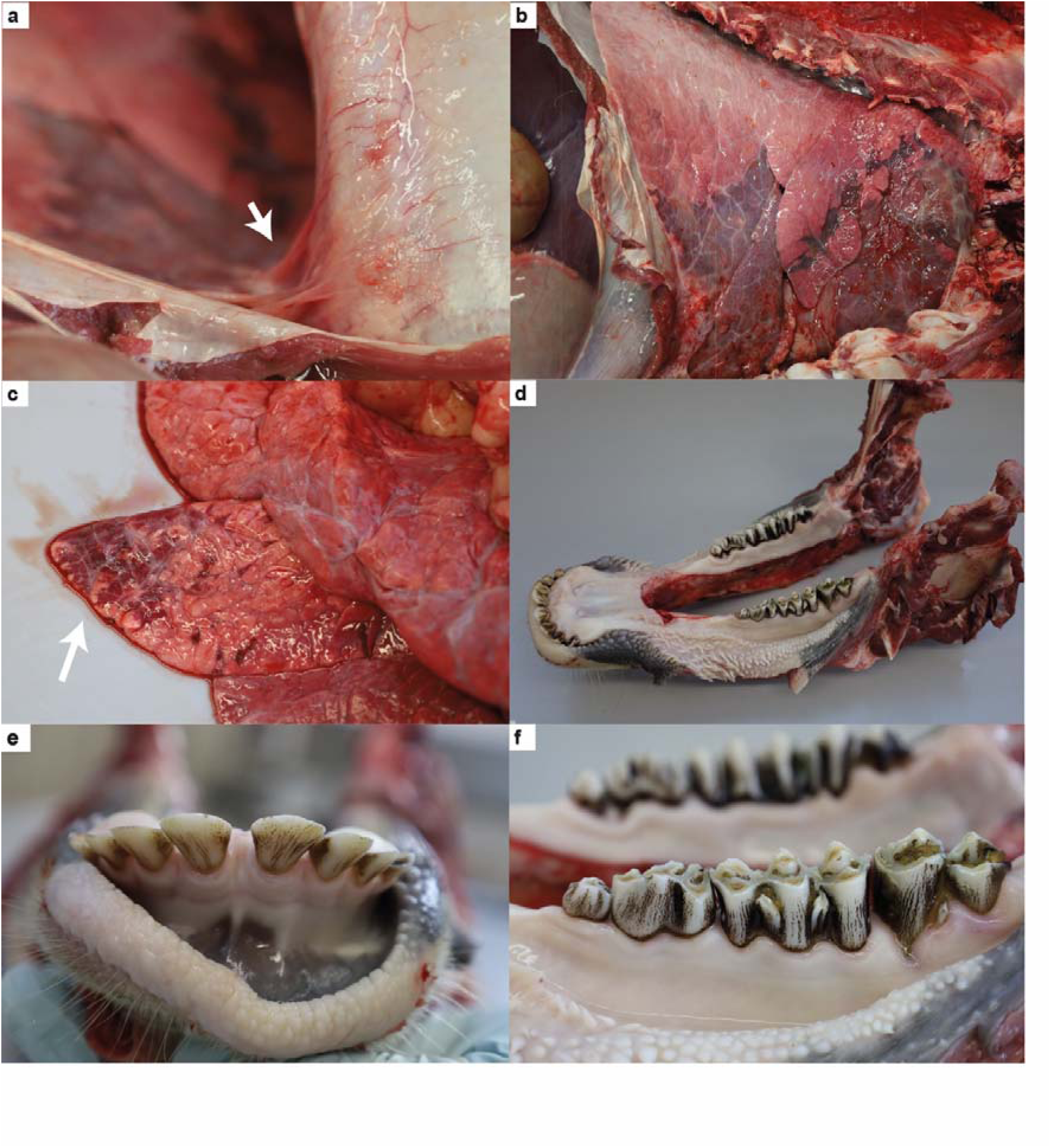
Pathological findings in BH39 homozygous Brown Swiss cattle. (a) Multifocal pleural adhesions (white arrow) connecting the lung to the costal pleura; the adhesions are band-shaped and fibrotic (case 1). (b) Cranial and middle lung lobes show dark red discoloration and increased consistency, consistent with bronchopneumonia (case 1). (c) Multifocal dark red discoloration (white arrow) with slightly increased consistency in the cranial lung lobe, consistent with bronchopneumonia (case 2). (d) Mandible with brown discoloration of the incisors, premolars and molars (case 2). (e) Multifocal brown discoloration of the incisors (case 1). (f) Multifocal brown discoloration of the premolars and molars (case 2).

Case 1 exhibited more severe and extensive lesions. The animal was markedly malnourished and had a rough, hypotrichotic hair coat. A subcutaneous interdigital ulceration (2 × 0.7 cm) was present on the right hindlimb. Within the thoracic cavity, multifocal fibrotic adhesions (Figure 3a) connected the right lateral lung surface to the costal pleura. Several ectatic bronchi were filled with purulent material.

Case 2 showed milder additional findings. The animal presented a reduced nutritional condition. The mucosa of the small intestine was mildly reddened. The external umbilicus was slightly enlarged and coarse, and both umbilical arteries were thickened and had not regressed appropriately for a 92-day old BS calf.

Histological examination of both cases revealed pulmonary and intestinal inflammatory changes. In both animals, the lungs showed multifocal inflammatory infiltrates within bronchi and alveoli, characterized by the presence of neutrophils and additional nucleated cells and occasionally accompanied by edema fluid, consistent with suppurative bronchopneumonia, which was more severe in case 1 than in case 2. Furthermore, both calves exhibited inflammatory alterations in the intestine with cellular infiltrates in the lamina propria. The jejunal mucosa contained round to oval, thick-walled coccidian developmental stages (∼150 µm) with numerous basophilic granules, consistent with *Eimeria* spp. Several intestinal crypts contained moderate numbers of partially degenerated neutrophils (crypt abscesses). Similar low-grade inflammatory infiltrates and crypt abscesses were present in the colon. Accordingly, both animals showed enteric inflammatory changes characterized by lymphoplasmacytic and eosinophilic infiltrates associated with crypt abscess formation. In addition, both animals exhibited multifocal to confluent brown discoloration of the teeth.

Case 1 had more severe and extensive lesions. These included a severe suppurative bronchopneumonia with multifocal abscess formation, marked interlobular edema, and multifocal fibrin deposits on the pleura. Moreover, the tracheobronchial lymph nodes exhibited moderate, chronic, purulent lymphadenitis with neutrophil-filled marginal sinuses and lymphocytic hyperplasia.

Case 2 showed additional mild changes. In the liver, multifocal periportal aggregates of small numbers of neutrophils, lymphocytes, and plasma cells were present, together with increased numbers of neutrophils in hepatic sinusoids. The liver also contained small, randomly distributed lymphocytic aggregates, and centrilobular hepatocytes showed few small, round, optically empty, intracytoplasmatic vacuoles consistent with lipid accumulation. In the kidneys, the renal medulla exhibited focal tubular accumulation of deeply basophilic granular material, consistent tubular mineralization.

Overall, the pathological findings were characterized by a consistent suppurative bronchopneumonia of variable severity and lymphoplasmacytic and eosinophilic enteritis with crypt abscess formation, accompanied by multifocal brown dental discoloration and lymph node enlargement, suggestive of an impaired ability to control inflammatory and infectious processes.

### Pedigree analysis supports recessive inheritance

A pedigree analysis was performed including the five clinical examined BH39 homozygous animals (cases 1-5). All animals could be traced back to a bull (red square) born in 1987, which represented the common ancestor connecting all five cases (Figure 4).

**Figure 4.**
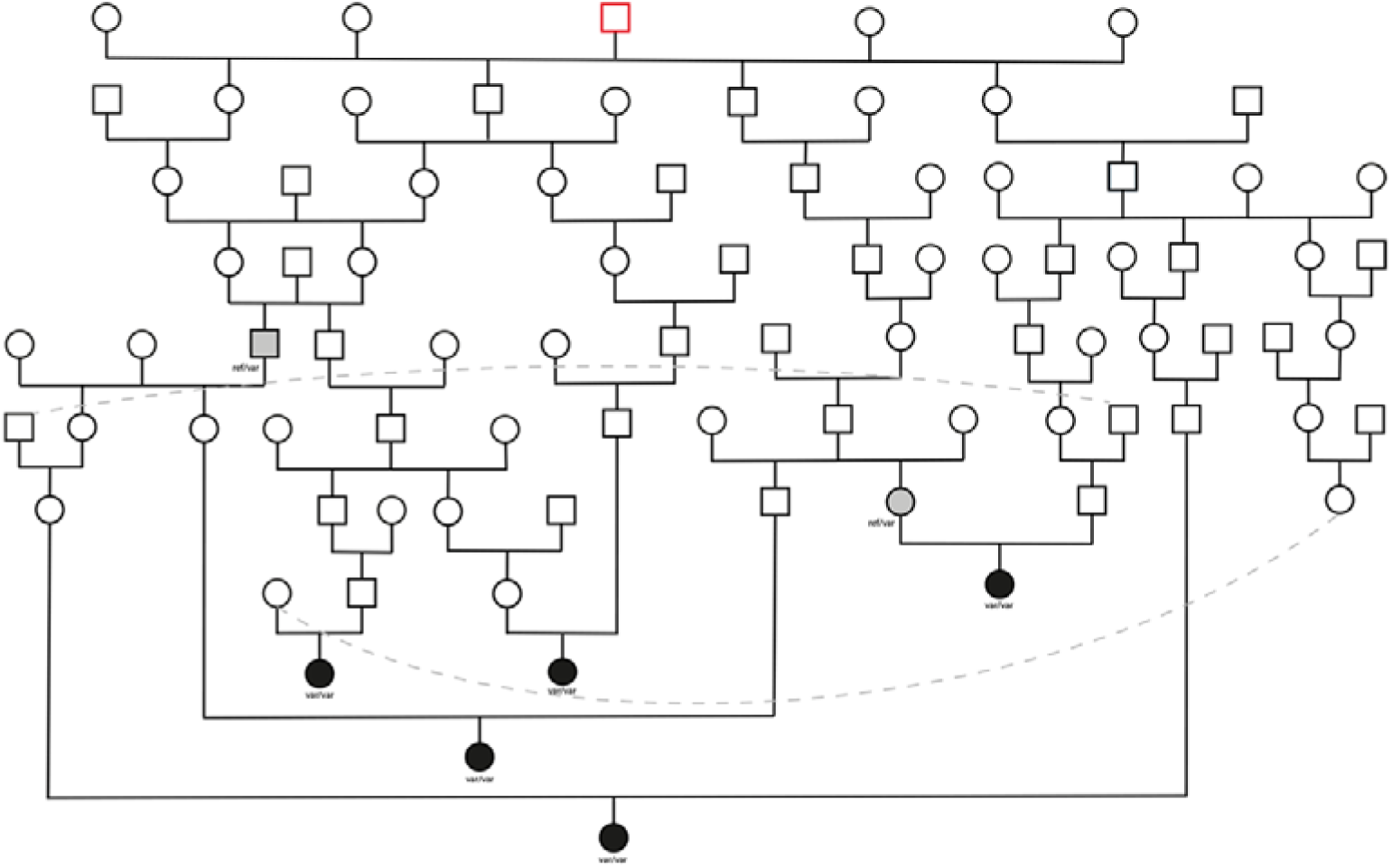
Pedigree analysis of the five clinical examined BH39 homozygous Brown Swiss cattle. Pedigree analysis revealed that five cases (black filled shapes) were related to a common ancestor indicating autosomal recessive mode of inheritance. Pedigree analyses indicate a bull born in 1987 (red-lined square) as common ancestor connecting all cases. Note the cross-generational matings and inbreeding loops leading to several individuals plotted repeatedly for the sake of readability (identical animals are shown as symbols connected with dashed lines). PCR-based *ALKBH8* genotypes (heterozygous: ref/var; homozygous: var/var) are presented for animals with DNA available.

Multiple inbreeding loops were identified in the pedigrees of the BH39 homozygous animals. The origin of the haplotype could not be conclusively determined.

### A 10.4 kb insertion of an ERV element in the coding sequencing of ALKBH8 is a causal variant

We analyzed imputed genotypes at 24,068,946 SNVs that were available for 58,871 BS animals to identify candidate causal variants for the increased juvenile mortality observed in homozygous haplotype carriers. Among these, 1541 SNVs on chr 15 were in high linkage disequilibrium (**LD**, r2 >0.7) with BH39. However, none of the variants in high LD were predicted to have a deleterious impact on protein function. Only three SNVs overlapped with the coding sequence, and all of them were annotated as synonymous variants (Supporting file S5). Thus, none of the SNVs represented a compelling candidate causal variant for the juvenile mortality of homozygous BH39 carriers.

Since the analysis of short-read sequencing-derived variants remained inconclusive, we performed HiFi sequencing on two BH39 homozygotes to investigate if the haplotype contains SVs that might explain the associated high juvenile mortality. Within a 10 Mb interval (15 – 25 Mb) on bovine chr 15, we identified 71 SVs that were homozygous for the alternate allele in both animals. Of these, 28 had an allele frequency of less than 10% (Supporting file S6) in a control group consisting of 84 animals of predominant BS ancestry. Only one of these variants overlapped with the coding sequence. This variant was called as a

10.4 kb insertion into the third exon of the *ALKBH8* gene at chr15:17045462 (Figure 5a-c). This insertion was not observed in the homozygous state in the 84 control animals. PCR-based genotyping confirmed that the other three clinically examined homozygous BH39 carriers were also homozygous carriers of the 10.4 kb insertion.

**Figure 5.**
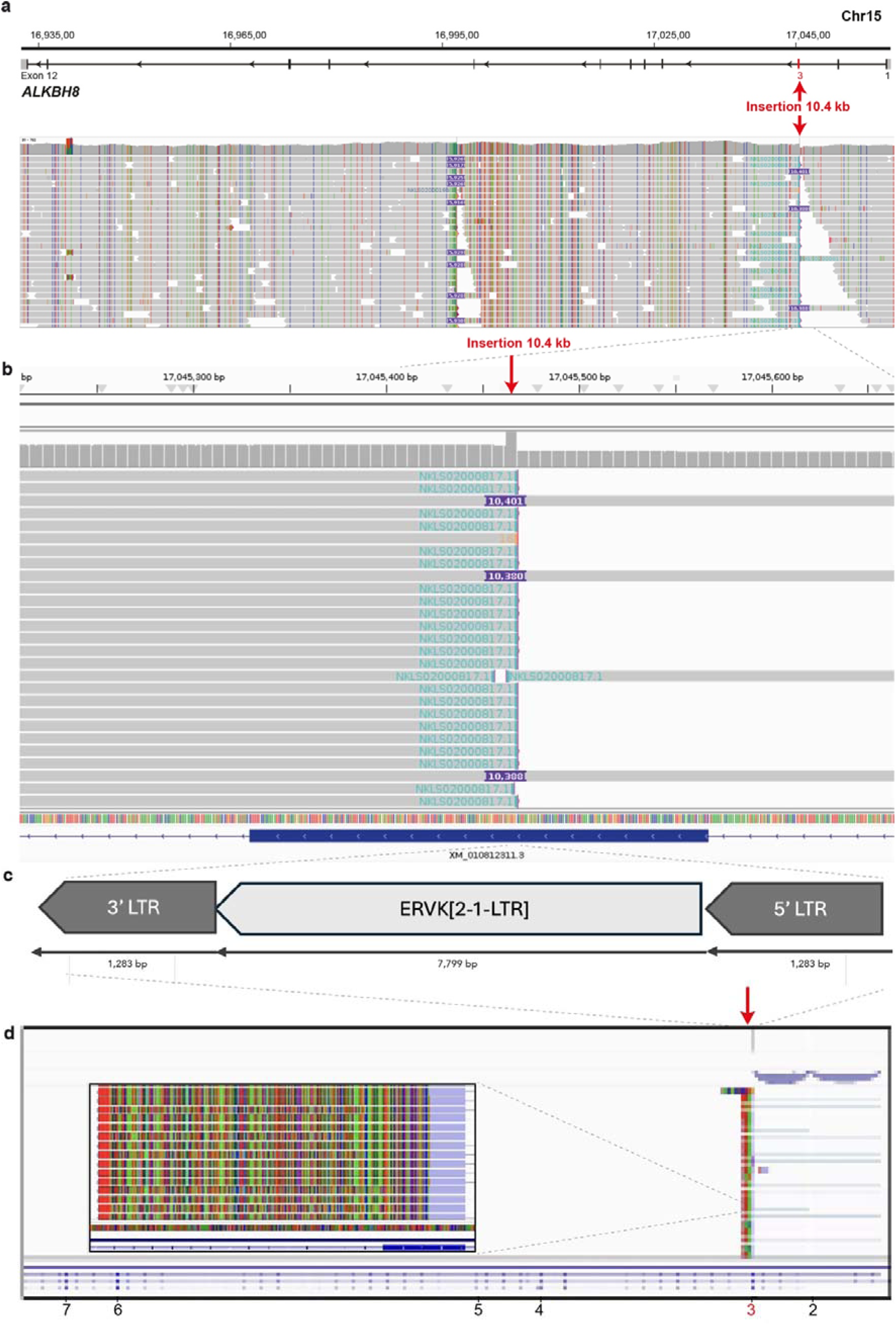
Insertion of an ERV element in the coding sequence of bovine *ALKBH8.* a) Structure of bovine *ALKBH8* gene on chromosome 15. The gene is transcribed from the reverse strand. A 10.4-kb insertion is located within exon 3 of *ALKBH8* (highlighted in red). b) IGV screenshots of PacBio HiFi read alignments from a homozygous BH39 carrier. Reads spanning the 10.4-kb insertion site show an excessive number of soft-clipped bases and increased read depth at the insertion locus. c) Magnified view of the genomic region shown in panel b, highlighting the 10.4-kb insertion site and the increased read coverage at the insertion site. d) Representation of the ERVK[2-1-LTR] element sense insertion and IGV screenshot of HiFi read alignments generated using PacBio Kinnex kits from liver RNA from a homozygous BH39 carrier showing a complete transcriptional shutdown in the ERV’s 5’LTR. The inset is a higher resolution visualization of a subset of the same alignments overlapping the third exon, showing soft-clipped based and the poly-A tail (red colour) marking the end of the transcript.

Analysis of the inserted sequence revealed that it is an ERV element that belongs to the [2-1-LTR] subgroup of ERVK elements. Visual inspection of the read HiFi alignments in IGV showed the expected abundance of both left-and right-clipped reads at the insertion site (Figure 5a-b). Moreover, a pronounced increase in read depth was observed across six nucleotides spanning positions 17,045,463 bp to 17,045,468 bp likely corresponding to the target site duplication (Figure 5b). The presence of a target site duplication is a key characteristic of ERV insertions and provides further support for the inferred insertion event.

The complete insertion is 10,404 bp long and consists of 7,799 bp sequence derived from the [2-1-LTR] subgroup of ERVK elements flanked by two 1,283 bp long terminal repeats (LTR, Figure 5c). Full-length transcript isoform sequencing using RNA prepared from cerebellum and liver tissue of a homozygous BH39 carrier showed that transcription is terminated in the 5’LTR of the ERV leading to a truncation of the *ALKBH8* open reading frame by 585 amino acids (87%, Figure 5d).

## DISCUSSION

This study applied a population-based genetic investigation that led to the discovery of a novel recessive deleterious haplotype in BS cattle (BH39) associated with rearing loss. By applying an HHED mapping approach, we identified a candidate recessive risk haplotype on chr15 that was associated, in homozygous state, with increased rearing loss within the first 180 days of life. Both, haplotype size and sliding window size, were fixed and did not vary in the HHED approach. This could have contributed significantly to the fact that the genomic segment of our candidate haplotype does not overlap well with the genomic position of the candidate causal variant identified subsequently. A French research consortium previously applied this strategy to uncover cryptic recessive disorders linked to increased mortality and premature culling rates in French cattle populations (Besnard et al., 2024).

To investigate the biological consequences of this haplotype, clinicopathological examinations were performed on BS homozygous haplotype carriers. These investigations revealed a multisystemic disorder primarily characterized by recurrent infections during early life and failure to thrive. Building on these findings, we focused on the candidate interval on chr15, where long-read whole-genome sequencing enabled the identification of a candidate causal structural variant in *ALKBH8*.

The identification of an inherited disorder with a clinical onset beyond the perinatal period highlights the methodological advantages of the HHED framework. Rather than exhibiting a single, distinctive cardinal sign of disease, homozygous risk haplotype carriers present with diffuse, multisystemic clinical manifestations, making recognition of the underlying genetic disorder based on clinical signs alone particularly challenging. As result, the unambiguous identification of affected and unaffected animals based only on the phenotype, which is a prerequisite for traditional case-control designs (Charlier et al., 2008), is not feasible. Our dataset comprised 891 animals that died before the age of 180 days of which only 10 (1.2%) were homozygous for BH39. Case-control association analysis with Fisher’s exact test of allelic association comparing animals that died within this age category to those who survived beyond this age would also identify the chr 15 region. However, the HHED approach provided a pronounced association signal demonstrating its utility to uncover cryptic recessive defects.

Following the identification of the associated genomic region, resolving the underlying genetic cause required a more comprehensive characterization of the candidate interval. Initial analysis of routine SNVs called from short-read sequencing did not reveal compelling list of candidate causal variants, leading us to hypothesize that the disorder could be caused by a structural variant. As short-read sequencing has inherent limitations in detecting and accurately genotyping SVs, PacBio HiFi sequencing was performed to enable comprehensive SV discovery and overcome these limitations (Logsdon et al., 2020; Nguyen et al., 2023). Long-read WGS of two BH39 homozygotes was therefore critical for resolving the genetic basis of the associated phenotype and identifying a candidate structural causal variant affecting *ALKBH8*. The variant consisted of a 10.4 kb endogenous retroviral insertion, highlighting the importance of interrogating repetitive and structurally complex regions that are often difficult to resolve with short-read WGS data alone. This finding is consistent with growing evidence that ERV mobilization remains an active source of mutational variation in cattle and can generate deleterious alleles (Tang et al., 2024). For example, an ERVK[2-1-LTR] insertion in exon 5 of *APOB* has been shown to cause inherited cholesterol deficiency in Holstein-Friesian cattle (Menzi et al., 2016). More broadly, recent work demonstrated that bovine ERVK[2-1-LTR] elements are still active in the germline, with mobilization rates varying substantially among individuals (Tang et al., 2024). That study further showed that competent C-type elements drive mobilization, whereas non-autonomous D-type elements can preferentially generate *de novo* insertions through trans-complementation and may act as “parasite-of-parasite” elements contributing to the long-term decline of ERV families. These observations support the interpretation that the BH39-associated insertion represents a plausible disease-causing structural variant and emphasize the value of long-read WGS for detecting, resolving, and annotating ERV-derived variants in cattle genomes (Nguyen et al., 2023; Grant et al., 2024; Tang et al., 2024).

*ALKBH8* encodes a tRNA-modifying enzyme essential for wobble uridine methylation and hydroxylation, ensuring accurate translational decoding (Van Den Born et al., 2011). Beyond its fundamental role in translation, *ALKBH8* appears to contribute to cellular stress responses, as loss of ALKBH8 function has been shown to reduce stress resilience and increase susceptibility to oxidative damage in mammalian models (Endres et al., 2015; Lee et al., 2020). In humans, biallelic variants in *ALKBH8* cause an autosomal recessive neurodevelopmental disorder (OMIM: 618504) further supporting the importance of *ALKBH8*-mediated molecular functions *in vivo* (Monies et al., 2019; Maddirevula et al., 2022; Yılmaz et al., 2024). Additional evidence suggests that the role of *ALKBH8* extends to the regulation of tissue responses to stress and inflammation. *Alkbh8*-deficient mice show increased susceptibility to oxidant-and toxicant-induced lung injury, indicating a broader role of *ALKBH8* in tissue stress responses (Leonardi et al., 2020). Moreover, in human sepsis, reduced *ALKBH8* expression has been associated with increased reactive oxygen species (**ROS**) production, activation of NFκB/NLRP3-mediated inflammatory pathways, and impaired cellular survival (Yang et al., 2021). Together, these findings suggest that ALKBH8 contributes to immune homeostasis during systemic infection and that reduced ALKBH8 activity may predispose to oxidative stress and dysregulated inflammatory responses. This biological context is consistent with the clinical phenotype observed in BH39 homozygous animals, in which recurrent respiratory disease and susceptibility to infections in some cases raised suspicion of an underlying immunodeficiency.

The marked brown discoloration of the teeth observed in our cases may reflect disturbed enamel development caused by oxidative stress secondary to repeated disease episodes. Although the precise pathogenesis remains uncertain, it is worth to highlight that enamel formation is highly dependent on proper ameloblast protein synthesis and secretion and appears vulnerable to disturbances in redox homeostasis, endoplasmic reticulum (**ER**) stress, and related cellular stress-response pathways (Sierant and Bartlett, 2012). In ameloblast cell models, experimentally increased oxidative stress elevates intracellular ROS, impairs mitochondrial bioenergetics, and reduces expression of enamel-specific genes critical for normal enamel formation (Li et al., 2022). Similarly, in a *Sod1*-knockout mouse model of oxidative stress, fluctuations in reactive oxygen species were associated with abnormal enamel structure and perturbation of regulatory pathways essential for amelogenesis (Xu et al., 2022). More broadly, enamel development can be disrupted by a variety of genetic and environmental stressors, including systemic infection, trauma, altered oxygen saturation, and certain drugs such as tetracycline (Xu et al., 2022; Wright, 2023). Further studies are needed to elucidate the precise mechanisms underlying this condition in BH39 homozygous cattle.

Beyond the dental phenotype, the clinical findings in BH39 homozygous animals indicate a broader impairment of resilience to physiological stress. Respiratory disease occurred frequently in BH39 homozygous animals, indicating decreased resilience of the lower respiratory tract, which is inherently susceptible to oxidative and inflammatory injury (Rogers and Cismowski, 2018; Bezerra et al., 2023). Hypoproteinemia might result from increased metabolic consumption during repeated disease episodes. Furthermore, reduced growth likely reflects inflammatory and metabolic stress. Further studies including a larger cohort of BH39 homozygous BS animals will be required to enable a more comprehensive phenotypic characterization, including detailed immunological profiling to assess the extent and nature of possible impairment of inflammatory response.

Interestingly, although the identified risk haplotype BH39 was associated with increased mortality during the rearing period, some homozygous animals survived beyond this stage. Notably, the oldest of the five clinically examined homozygous animals showed no remarkable clinical abnormalities apart from anestrus and dental abnormalities. This observation raises the possibility of incomplete penetrance and/or variable expressivity of the phenotype. Such variability may reflect the influence of environmental factors, management conditions, or stochastic biological processes that modulate individually different disease severity (Kingdom and Wright, 2022). However, given the limited number of homozygous animals available for clinical evaluation, no firm conclusions can be drawn. Future studies including larger cohorts and longitudinal phenotypic characterization will be needed to determine the extent of incomplete penetrance and the factors contributing to phenotypic variability.

Taken together, these findings contribute to a growing understanding of BH39 as a complex recessive disorder with variable clinical expression and important implications for cattle health. Our study aligns with recent advances in genomic selection programs that increasingly integrate causal variants to reduce juvenile losses in dairy populations (Chapard et al., 2025). Early identification of carriers and informed mating decisions will be essential to prevent additional cases and improve animal welfare.

Several limitations should be considered when interpreting the phenotypic characterization of BH39 homozygous animals. The number of available homozygous animals for detailed clinical examination was limited, and animals were included based on survival to the time of examination and owner consent. Therefore, ascertainment bias cannot be excluded, as the examined animals may not fully represent the spectrum of disease severity, particularly the most severe early-life manifestations. In addition, the HHED control definition, which required animals to have produced offspring, may introduce a selection bias by enriching controls for animals with adequate health and reproductive performance.

## CONCLUSIONS

The clinical, pathological, and genetic findings provide strong evidence that the BH39-related *ALKBH8* loss-of-function variant described here causes a new syndromic recessively inherited Mendelian disorder in BS cattle. Our study demonstrates how analyzing life trajectories and big data driven genomics enables proactive observation of hidden harmful recessive alleles. Early identification of affected animals and carriers is not only essential for breeding management, but also has important implications for animal welfare, as homozygous animals are frequently affected by recurrent disease and require repeated therapeutic interventions. Preventing the birth of affected calves may therefore reduce animal suffering and contribute to a decrease in antibiotic use associated with repeated treatments. Further molecular and phenotypic investigations will be required to clarify the pathogenic mechanisms and to fully assess the phenotypic spectrum of this disorder.

## SUPPLEMENTARY MATERIAL

**File S1**

Format: .csv

Title: Diagnostic test 15_017045468 *ALKBH8*.

**File S2**

Format: .docx

Title: Survival analysis of Swiss, German and Austrian BS cattle populations.

**File S3**

Format: .xlsx

Title: Signalment and clinical findings in five BH39 homozygous animals.

**File S4**

Format: .xlsx

Title: Hematology and blood biochemistry results in five BH39 homozygous animals.

**File S5**

Format: .vep

Title: Variant Effect Predictor

**File S6**

Format:.txt

Title: BH39_28SVs

Reserved DOI: 10.17632/trfjggcyyp.1

## NOTES

### Ethics approval and consent to participate

All animals in this study were examined with the consent of their owners and handled in accordance with the Swiss Animal Welfare Act and institutional ethical guidelines (Swiss Federal Food Safety and Veterinary Office, 2024). The SNV array genotypes were generated during population-wide genotyping of Swiss dairy populations for the purpose of genomic selection. Collection of blood samples was approved by the Cantonal Committee for Animal Experiments (Canton of Bern; permit BE94/2022).

### Author contributions

The author contributions were as follows: Sidonia Glatthard writing (review and editing), methodology, investigation, formal analysis, data curation. Naveen K. Kadri, writing (review and editing), methodology, investigation, formal analysis, software. Franz R. Seefried, writing (review and editing), methodology, investigation, formal analysis, conceptualization, funding acquisition, supervision, project administration. Laura R. Voitl, writing (review and editing), methodology, investigation, formal analysis. Bettina A. Weber, investigation, formal analysis. Hermann Schwarzenbacher, investigation, formal analysis. Seraina L. Meister, methodology, investigation, formal analysis. Corinne Gurtner, methodology, investigation, formal analysis. John F. O’Grady, methodology, investigation, formal analysis. Meret Osbahr, methodology, investigation, formal analysis. Alexander S. Leonard, methodology, investigation, formal analysis. Mireille Meylan, writing (review and editing), methodology, investigation, formal analysis. Hubert Pausch, writing (review and editing), conceptualization, funding acquisition, supervision, project administration. Cord Drögemüller, writing (review and editing), conceptualization, funding acquisition, supervision, project administration. Joana Jacinto, writing (review and editing), methodology, investigation, formal analysis, conceptualization, funding acquisition, supervision, project administration.

## Acknowledgements

The authors acknowledge Isabella Aebi-Huber and Edoardo Henzen for expert technical assistance. We thank the Next-Generation Sequencing Platform of the University of Bern and the Functional Genomics Center Zurich for performing the hight-throughput sequencing experiments and Interfaculty Bioinformatics Unit of the University of Bern for providing a high-performance computing infrastructure. We would also like to thank Braunvieh Schweiz, Swissgenetics, Select Star SA, and all the owners who contributed samples and information about cases.

