## Supporting File S1 for "An endogenous retrovirus insertion disrupting bovine *ALKBH8* causes a failure-to-thrive syndrome with immunodeficiency associated with juvenile mortality in Brown Swiss cattle"

**Supporting File 1: Survival analysis of Swiss, German and Austrian BS cattle populations.**

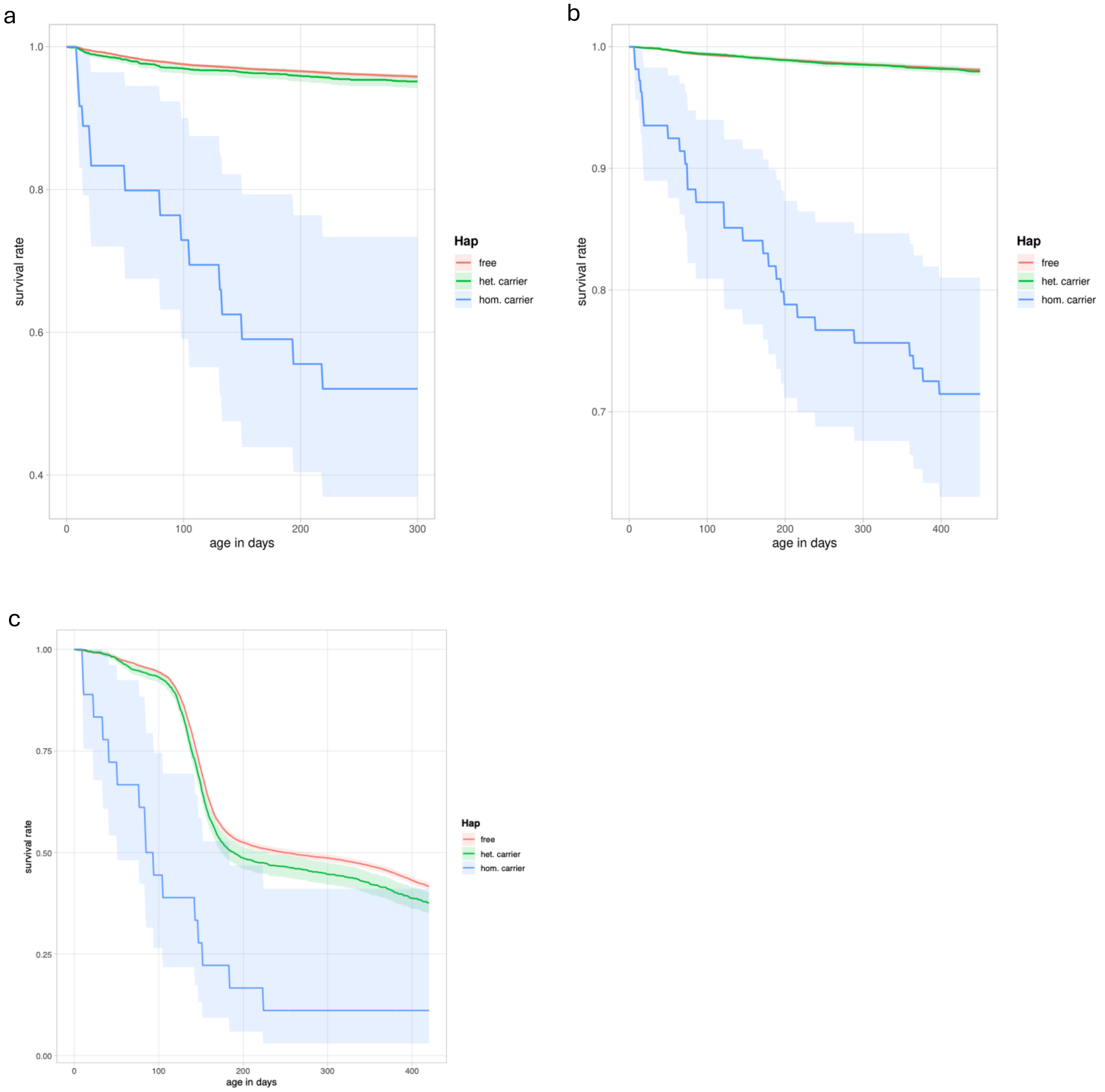


Male (a) and female (b) survival rate estimated in 127,055 BSW cattle in the German/Austrian population. The blue line represents the survival rate of homozygous haplotype carriers. Blue shaded area is the confidence interval. Survival rate of heterozygous carriers (green) is indistinguishable from the survival rate of non-carriers (red). Age at genotyping differs significantly between populations. In Austrian and German BS population, 50% of females are genotyped later than 166 days of age, whereas in the Swiss BS population animals are genotyped much earlier, with a median of 65 days in recent cohorts. In the Swiss male population (c), a marked decline in survival is observed around 160 days of age in heterozygous carriers (green) and non-carrier animals (red). This decrease can be explained by the Swiss agricultural production system, where calves intended for veal production are typically slaughtered at this age. In contrast, this decline around 160 days is absent in the German/Austrian population, likely reflecting differences in production systems between the countries.
